# In vitro comparison of the internal ribosomal entry site activity from rodent hepacivirus and pegivirus

**DOI:** 10.1101/761379

**Authors:** Stuart Sims, Kevin Michaelsen, Sara Burkhard, Cornel Fraefel

**Affiliations:** Institute of Virology, University of Zurich, Zurich, Switzerland; Clinic for Internal Medicine, University Hospital of Zurich, Zurich, Switzerland

**Keywords:** Hepacivirus, IRES, HCV, Pegivirus

## Abstract

The 5’ untranslated region (5’ UTR) of rodent hepacivirus (RHV) and pegivirus (RPgV) contain sequence homology to the HCV type IV internal ribosome entry sites (IRES). Utilizing a monocistronic expression vector with an RNA polymerase I promoter to drive transcription we show cell-specific IRES translation and regions within the IRES required for full functionality. Focusing on RHV we further pseudotyped lentivirus with RHV and showed cell surface expression of the envelope proteins and transduction of murine hepatocytes.

## Introduction

Hepatitis C virus (HCV) infects more than 71 million people worldwide(1) leading to liver failure and hepatocellular carcinoma, presenting a major global health burden. While the introduction of new direct-acting antiviral drugs (DAAs) has improved treatment response rates and heralded a new era of HCV treatment(2), the cost and availability of DAAs along with drug resistance, and chronic/non-reversible liver damage due to late onset of symptoms and delayed treatment initiation after HCV infection are an issue(3). A small animal model for HCV would allow for the testing of vaccines which could potentially solve these problems.

HCV was discovered in humans 20 years ago(4) but for a long-time researchers failed to identify an animal viral homologue, this all changed with the use of high throughput deep sequencing and has shed light on the evolutionary origins of the virus. The first HCV homologue was discovered in 2011 in canines(5) and was quickly followed by the discovery of homologues in rodents(6,7), equine(8,9), primates(10), bovine(11,12) and then, the first nonmammalian species, sharks(13).

The rodent HCV homologues were termed rodent hepaciviruses (RHV) and were first identified in deer mice (*Peromyscus maniculatus*), a species known to carry hantavirus, desert woodrats (*Neotoma lepida*) and hispid pocket mice (*Chaetodipus hispidus*)(7). The same year RHV was described in European bank voles (*Myodes glareolus*) and South African four-striped mice (*Rhabdomys pumilio*)(6). This was succeeded by the discovery of an RHV from Norway rats (*Rattus norvegicus*) in New York City(14).

The subsequently discovered RHV (RHV-rn1) from Norway rats was used to make a small animal model for hepaciviral infection, utilising a reverse genetics approach. In this system researchers showed its hepatotropic replication in inbred and outbred rat strains(15). Emulating HCV infection they also showed that persistent infection leads to gradual liver damage and that the HCV antiviral drug sofosbuvir suppresses replication of RHV-rn1. This model can be used to study the mechanisms of HCV persistence, immunity, and pathogenesis.

Though rats are the usual hosts of RHV-rn1, it has also been shown that the virus is capable of establishing a persistent infection in immunocompromised mice lacking type I interferon and adaptive immunity(16). However, immunocompetent mice clear the virus in a few weeks. Because this mouse model only results in an acute infection, a fully immunocompetent mouse model in which a chronic infection and downstream liver damage can be established is still in need. Additionally, the availability of knock out mice would aid the study of the pathogenesis of HCV related liver damage.

While these studies could lead the way to future vaccines they also present the possibility of zoonotic sources of HCV infection in humans(17). The genomes of RHV encode for a polyprotein, that is predicted to be cleaved into 10 proteins, as with HCV, but shows a 66-77% amino acid divergence from HCV in the structural genes(18),(7), therefore, tropism and pathogenicity may also differ between the viruses. In particular, the 5’ untranslated regions (UTRs) of RHV are highly divergent from the corresponding regions of HCV and there is a large difference between the different rodent clades (RHV, RHV1, RHV2, RHV3, RHV-rn1).

The high-level expression of mir-122 in liver cells allows HCV, containing two mir-122 target sequences in the 5’UTR, to replicate and is one reason for HCV hepatotrophism(19). Similarly, the RHV (NC_021153) found in deer mice contains one such mir-122 target sequence in its 5’UTR(7) suggesting liver cell specificity and further has a 200nt sequence with no homology to other hepaciviruses 5’UTRs. The RHV-1 (KC411777)) 5’UTR contains structural elements typical of both pegi- and HCV-like IRES and contains one mir-122 target region. While RHV-1 and RHV-2 (KC411784) are identical in structure and only contain a few nucleotide exchanges, with RHV-3 (KC411807) and RHV-rn1 being more similar to typical HCV-like IRES structures(6,15).

Along with RHV, a rodent pegivirus (RPgV) was also discovered in white-throated wood rats (*Neotoma albigula*)(7). Pegiviruses are a new genus of the family of the Flaviviridae, encompassing the human GBV-A, GBV-C/HGV/HPgV-1, GBV-D and HPgV-2 viruses. These pegiviruses are considered to be non-pathogenic and in the case of HPgV has even been reported to be beneficial in co-infections with HIV or Ebola(20–22). However, two pegiviruses were discovered in equine, the first, Theiler’s disease-associated virus (TDAV) is suspected to be the causative agent for an outbreak of acute hepatic disease occurring on a horse farm(23). The second, equine pegivirus (EPgV), like the human pegiviruses, is considered to be non-pathogenic(24).

The viral RNA structures play important roles in both translation and replication. Specifically, the 5’UTR containing the IRES promotes the initiation of protein synthesis in a cap-independent manner. IRES’s are diverse in sequence and structure and these differences contribute to tropism; the focus of this study is to assess how these differences in the 5’UTRs of RHV and RPgV affect translation and to establish a reverse genetics system.

## Materials and Methods

### Cells

Hepa1-6 (ATCC^®^ CRL-1830™), MEFs IRF3^−/−^, NIH 3T3 (ATCC^®^ CRL-1658™), BHK-21 (ATCC^®^ CCL-10™), HEK 293T (ATCC^®^ CRL-11268™), Huh-7.5-RFP-MAVS from Dr Charles M Rice, HepG2 (ATCC^®^ HB-8065™) and Vero cells (ATCC^®^ CCL-81™) were cultured in DMEM (Sigma-Aldrich), 10% FBS and 1% penicillin-streptomycin.

### Plasmid construction

Monocistronic reporter plasmids containing viral 5’UTR were constructed in pUC19. First pUC19 was digested with EcoR1 and HIndIII (NEB), then a minimal RNA polymerase I (RNA Pol I) promoter and terminator (synthesised by Twist Biosciences) was inserted into the digested plasmid by In-Fusion cloning (Takara Bio), this plasmid was named pOLI (Fig S1A).

The plasmid pOLI was linearized with PpuMI. Viral 5’ and 3” UTR (synthesised by Twist Biosciences) (HCV IRES taken from pFR_HCV_xb (Addgene)) along mCitrine were cloned into the linearized pOLI by In-Fusion cloning (Takara Bio) (Fig S1B). Deletions and additions to the monocistronic reporter plasmids were created by PCR and In-Fusion cloning (Takara Bio). Plasmids containing viral structural genes were constructed in pUC19 by In-Fusion cloning (Takara Bio) using hepacivirus CE1E2 region (synthesised by Twist Biosciences), along with a CMV promoter and BGH polyA (Fig S1C). Flag tag and c-Myc tag sequences were inserted into this plasmid by PCR and In-Fusion cloning (Takara Bio), this plasmid was named pC^Myc^E1E2^Flag^. The capsid gene was deleted from pC^Myc^E1E2^Flag^ by PCR and In-Fusion cloning (Takara Bio) resulting in the plasmid pE1E2^Flag^.

To construct plasmids containing full-length RHV1 and RPgV viral genomes, the respective monocistronic vector (pOLI.IRES.Cirtine) were PCR linearized and gene fragments, 1.5-2kb (synthesised by Twist Biosciences), covering the full-length viral coding region were then cloned in by In-Fusion cloning (Takara Bio) replacing mCitrine. The plasmids were named pOLI.RHV1 and pOLI.RPGV (Fig S1D).

For constructing a reporter plasmid, mScarlet -BSD (synthesised by Twist Biosciences) was cloned into the plasmid containing the full-length virus in-between NS5A and NS5B while also duplicating the cleavage sequence by In-Fusion cloning (Takara Bio). These plasmids were named pOLI.RHV1.SB and pOLI.RPgV.SB (Fig S1E).

### Transfection

All transfections were carried out using Lipofectamine^®^ LTX with Plus™ Reagent (Thermo Fisher Scientific) in Opti-MEM (Gibco) according to the manufacturer’s specifications.

### Western blot

Cells were lysed in RIPA buffer (150 mM NaCl, 50 mM Tris/HCl pH 7.6, 1% Nonidet P40, 0.5% sodium deoxycholate, 5 mM EDTA) supplemented with cOmplete™ Protease Inhibitor Cocktail (Roche) for 15 min on ice. The lysate was run on a 12% SDS-Page and transferred to PVDF membrane (Bio-Rad). Membranes were incubated for 1 hour in PBS with 5% non-fat dry milk, then stained with primary and subsequently secondary antibodies for 1 hour in PBS with 1% non-fat dry milk. Immunocomplexes were detected using an Odyssey^®^ Fc Imaging System (LI-COR Biosciences).

### Immunofluorescence

#### Intracellular

Cells were grown on glass coverslips, washed with PBS prior to fixation in formaldehyde (3.7% w/v in PBS) for 10 min at room temperature, permeabilized for 5 min with 0.1% Triton X-100 in PBS, and blocked with 3% BSA in PBS for 30 min. Primary and secondary antibodies were diluted in PBS containing 1% BSA, cells were stained with primary antibody for 1 hour, washed 3x with PBS then stained with secondary antibody for 1 hour, followed by staining with 0.1 μg/ml DAPI in PBS for 5 min and application of Prolong Gold Antifade reagent (Invitrogen).

#### Extracellular

Cells were grown on glass coverslips, incubated with PBS containing 4% FBS (Fetal Bovine Serum) for 30min. Primary and secondary antibodies were diluted in PBS containing 4% FBS, cells were stained with primary antibody for 1 hour, washed 3x with PBS then stained with secondary antibody for 1 hour. Cells were then stained with Wheat Germ Agglutinin, Alexa Fluor™ 594 conjugate as per manufacturers protocol. Cells were fixed in formaldehyde (3.7% w/v in PBS) for 10 min at room temperature, permeabilized for 5 min with 0.1% Triton X-100 in PBS and stained with 0.1 μg/ml DAPI in PBS for 5 min followed by application of Prolong Gold Antifade reagent (Invitrogen).

Images were acquired using a confocal microscope (Leica specify type), Z-stacks, and analysed with ImageJ.

### Antibodies

Rat anti-DYKDDDDK (clone L5, Biolegend), mouse anti-c-myc (clone 9E11, Biolegend), mouse anti-β-actin (clone 2F1-1, Biolegend), mouse J2 anti-dsRNA IgG2a (Sciscons), goat anti-rat IgG (H+L) Alexa Fluor 488 (Invitrogen), IRDye^®^ 800CW goat anti-rat IgG and IRDye^®^ 680RD donkey anti-mouse IgG (LI-COR Biosciences).

### Flow cytometry

Single cell suspensions were generated and kept in FACS buffer (2% FCS, 5mM EDTA in PBS). Cells were analyzed using a Gallios flow cytometer (Beckman Coulter) and FlowJo software, and gated on viable cells using the live/dead fixable near-IR dead cell stain kit (Invitrogen).

### Lentivirus

Plates were seeded with HEK 293T in DMEM and 3% FCS, then transfected with pLKO-gfp (Addgene), pCMVΔR8.2 (Addgene) and either pMD2.G (Addgene) or pE1E2 at a ratio of 1:1:0.1 respectively using lipofectamine LTK (Thermo Fisher Scientific) and Opti-mem (Gibco). At 6 hours post transfection, media was replaced and 72h post transfection, supernatant was harvested and passed through a 0.45μm filter.

### GeneBank accession numbers

RHV (Hepacivirus E); NC_021153, RHV1 (Hepacivirus J); KC411777, RHV2 (Hepacivirus F); KC411784, RHV3 (Hepacivirus I); KC411807, RHV-rn1 (Hepacivirus G); KX905133.1, RPgV; NC_021154.

## Results

### RHV and RPgV IRES’s are functional in Rodent cells

To test viral IRES driven translation, a monocistronic plasmid vector was constructed containing a minimal RNA polymerase I promoter in front of the full-length viral 5’ UTR followed by a fluorescent maker, the viral 3’ UTR, and the RNA polymerase I terminator (Fig 1A). The HCV IRES was used as a positive control along with a control plasmid containing a random sequence in place of the viral 5’ UTR. We constructed plasmids containing RHV, RHV1, RHV2, RHV3, RHV-rn1 and RPgV 5’ UTR from previously published sequences.

**Fig1.**
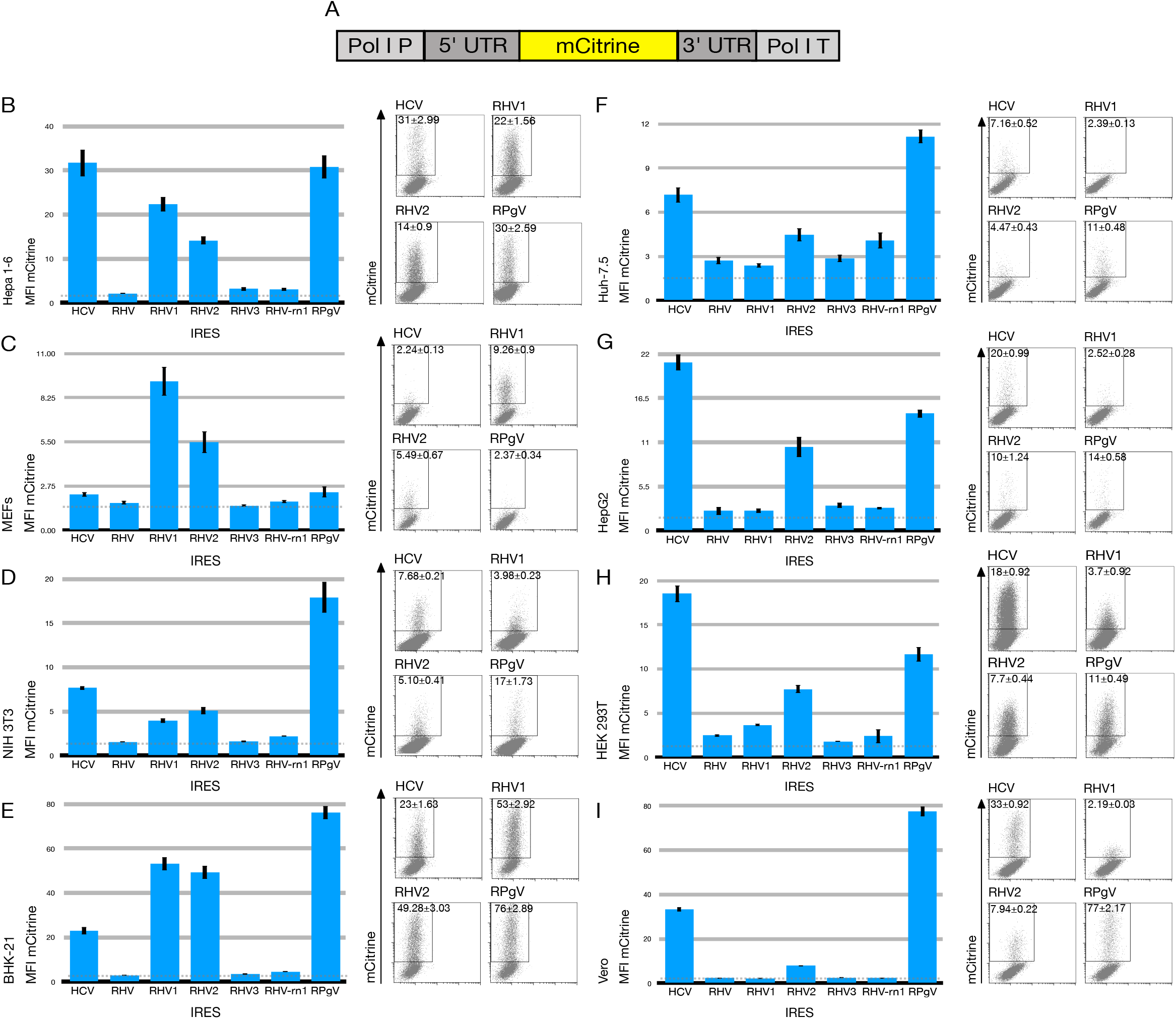
RHV and RPGV IRES activity in different cell types. Schematic of the monocistronic vectors used (A). Hepa1-6 (B), MEFs (C) NIH 3T3 (D), BHK-21 (E), Huh7.5 (F), HEK 293T (H) and Vero (I) cells were transfected with the indicated plasmids. Cells were harvested and analysed by flow cytometry at 48 h.p.t, bar graphs show MFI for mCitrine (n≥8, mean±SEM of at least three independent experiments) and representative FACS plots on the right of each graph.

These plasmids were transfected into the murine hepatocyte cell line Hepa1-6, with the HCV and RPgV IRES driving the highest level of translation at 72 hours post transfection with a mean fluorescence intensity (MFI) of 30, followed by RHV1 and RHV2 with an MFI of 22 and 14, respectively. RHV3 and RHV-rn1 IRES drove translation at a level only slightly above background and RHV was not functional in Hepa1-6 (Fig 1.B).

To further asses IRES function in murine cells, we tested two murine embryonic fibroblasts cell lines, MEFs and NIH 3T3. Transfection of MEFs with the plasmids revealed that RHV1 drove the highest level of translation at an MFI of 9 followed by RHV2 at an MFI of 5 (Fig 1.C). HCV, RHV, RHv3, RHV-rn1 and RPgV generated signals only slightly above the negative control. In NIH3T3 the RPgV drove the highest level of translation with an MFI of 17 followed by HCV with an MFI of 7, again RH1 and RHV2 were functional but at low levels and RHV, RHV3 and RHV-rn1 were not functional (Fig 1.D).

The viruses originate from different rodent species therefore the baby hamster kidney cell line was tested to assess if the IRES’s are functional in this cell line. As with the previous cell lines, the RPgV IRES drove high levels of translation, also the RHV1 and RHV2 were capable of driving high levels followed by HCV, again the RHV, RHV3, RHVrn1 were not functional (Fig 1.E).

### RHV and RPgV IRES’s show differing function in human cell lines

In the human hepatocyte cell line Huh7.5 which expresses high levels of mir122, the RPgV IRES drives the highest levels of translation followed by HCV and RHV2. The RHV-rn1 drove low levels of translation and, as with murine cells, RHV and RHV3 did not yield any signal (Fig 1.F). Another human hepatocyte cell line, HepG2 that does not express mir122, was also used to test the IRES’s function: in these cells, HCV drove the highest levels followed by RPGV and RHV2, while the expression from RHV, RHV1, RHV3 or RHV-rn1 were at background level (Fig 1.G).

To investigate if the IRESs are functional in non-hepatocyte cells lines, we used HEK 293T, and Vero cell lines. In HEK 293T the HCV IRES showed the highest level of translation followed by the RPgV IRES and RHV2 while that of RHV, RHV1, RHV3 and RHV-rn1 were at the lowest level (Fig 1.H). In Vero cells The RPgV produced high levels of translation, over twice that of HCV; RHV2 was the only other IRES that was functional in these cells although at a very low level when compared to RPGV (Fig 1.I).

### Deletions abrogate the function of RHV1 and RPgV IRES

To further asses the structural requirements of RHV1 IRES for full functionality we made several different constructs. The first contained an additional 20 nucleotides of virus sequences downstream of the start codon. When transfected into Hepa1-6 cells this construct did not lead to a difference in the levels of translation in comparison to the construct containing just the 5’ UTR. Two deletion constructs were made, in RHVΔI the 5’ three stem loops (Ia/b/c) were deleted, this deletion decreased the IRES function by 90%. The Va and Vb stem loops were deleted from RHV1ΔII and again led to a decrease in function by 90% in the murine hepatocyte cell line Hepa1-6 (Fig 2.C).

**Fig2.**
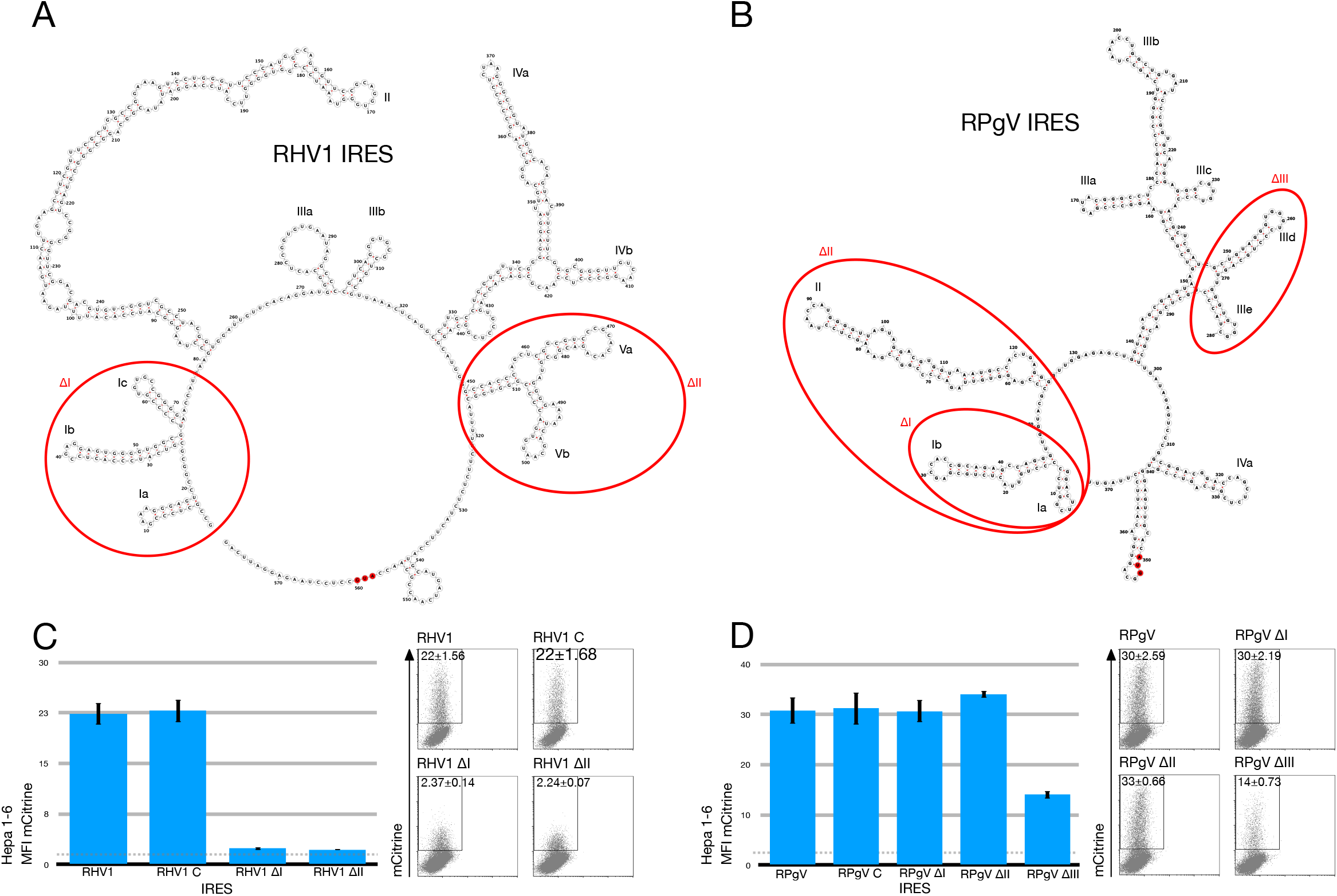
Deletions to RHV1 and RPgV 5’ UTR abrogate IRES translation. Predicted RNA secondary structure of RHV1 (A) and RPgV (B). Red circles indicate areas deleted from plasmids. Hepa1-6 cells were transfected with RHV1 (C) and RPgV (D) plasmids. Cells were harvested and analysed by flow cytometry at 48 h.p.t. Bar graphs show MFI for mCitrine (n≥8, mean±SEM of at least three independent experiments) and representative FACS plots on the right of each graph.

For the RPgV IRES we also added an additional 20 nucleotides of virus sequences downstream of the start codon. This again had no effect on the levels of translation in comparison to the construct containing just the 5’ UTR. To further asses the sequence required for driving translation, three constructs were made with deletions, RPgVΔI and RPgVΔII have deletions to the 5’ of the IRES with RPgVΔI having the first two stem loops (Ia/b) and RPgVΔII three stem loops deleted (Ia/b and II). RPgVΔIII has two internal stem loops deleted (IIId/e). The 5’ deletions to RPGV had no effect on the levels of translation, however the internal deletions in RPgVΔIII led to a reduction in translation of 53% (Fig 2.D).

### Expression of E1E2

Previous studies examining the subcellular localization of HCV E1 and E2 used cells transfected with a plasmid expressing the E1 and E2 proteins(25). These studies concluded that the HCV structural proteins are expressed on the cell surface, based on immunofluorescence detection. In order to examine the localization of RHV structural proteins in an expression system we cloned the structural region (capsid-E1-E2) of RHV into a plasmid containing the CMV promoter, we then added the c-Myc tag to the 5’ end of the capsid protein and the Flag tag to the 5’ end of the E2 protein following the E1E2 cleavage sequence (Fig3.A).

**Fig3.**
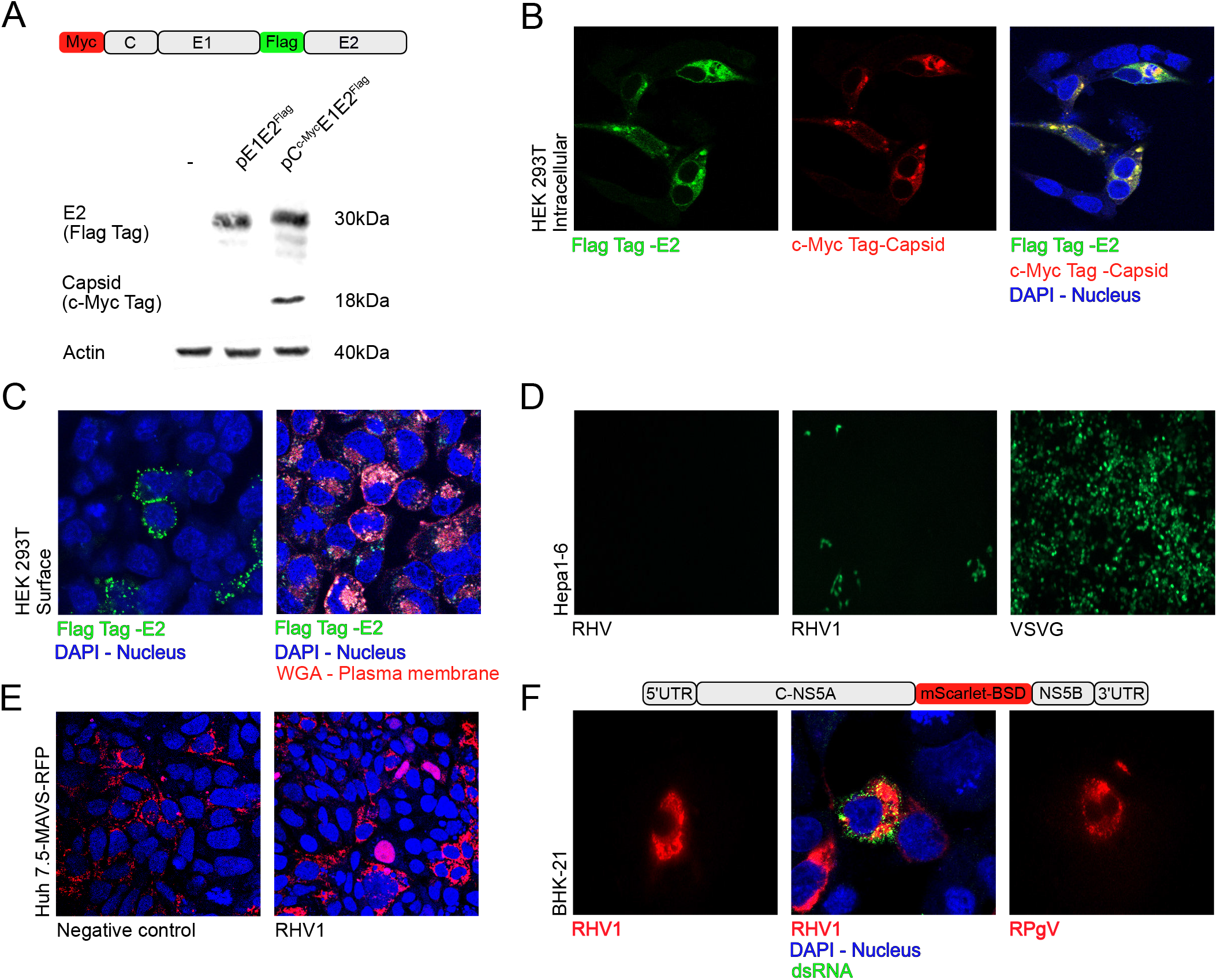
Expression of RHC envelope proteins and full-length recombinant virus. Schematic of the RHV1 envelope expression vector and Western blot of cell lysate at 48h.p.t from transfected HEK 293T (A). Intracellular immunofluorescence of 293T cells at 48h.p.t with RHV1 envelope expression vector, stained with flag tag in green, c-Myc tag in red and combined with DAPI stain (B). Extracellular immunofluorescence for E2 by Flag tag stain (green) and combined with WGA (Red) and DAPI (Blue) from 293T at 48h.p.t with RHV1 envelope expression vector, images represent one slice from z-stack (C). GFP-lentivirus vector pseudotyped with RHV or RHV1 envelope proteins or VSVG were incubated on Hepa1-6 cells; GFP positive cells indicate transduction (D). Huh7.5-MAVS-RFP cells transfected with full length RHV1 expression plasmid. Translocation of RFP to the nucleus is indicated by arrows (E). BHK-21 cells transfected with full length recombinant RHV1 or RPGV expression plasmid containing mScarlet (Red) as in schematic, combined with dsRNA staining (Green) and DAPI (Blue) (F).

HEK 293T cells were transfected with the vectors pE1E2^Flag^ or pC^Myc^E1E2^Flag^ and after 48 hours lysed for SDS-PAGE. The E2 protein was detected in cells transfected with either expression vector using an anti-Flag tag antibody (Fig3.A), and the capsid was detected in cells transfected with the pC^Myc^E1E2^Flag^ expression vector using an anti-c-Myc antibody. The proteins detected were of the predicted size, showing post-translational cleavage was complete.

In order to determine if the RHV glycoproteins expressed from these vectors also exhibit an intracellular colocalization, we examined transfected cells by immunofluorescence for capsid and E2 expression; both were shown to colocalize (Fig3.B). We also assessed if the envelope protein was expressed on the cell surface. Staining with anti-flag antibody to detect E2 indeed showed punctate staining on the cell membrane and when combined with wheat germ agglutinin to stain the cell membrane it showed colocalization with the flag antibody (Fig3.C). This indicates that a proportion of E2 is surface-expressed.

To determine if the surface-localized E1E2 could mediate viral entry, we produced a GFP encoding lentivirus vector pseudotyped with the RHV, RHV1, RHV2 and RHV3 E1E2 proteins by transfecting HEK 293T cells with pE1E2 and the lentivirus backbone and packaging plasmid, and 72h later harvesting and filtering the supernatant. We then tested if the E1E2-pseudotyped lentivirus vectors were entry-competent. Supernatants from the co-transfected cells were applied to Hepa1-6 cells and GFP reporter expression assayed 72 h later. RHV1 E1E2-pseudotyped lentivirus vectors gave rise to a small number of GFP positive cells, when compared to VSVG pseudotyped lentivirus. Lentivirus vectors pseudotyped with RHV, RHV2, RHV3 or lacking envelope glycoprotein failed to give rise to any GFP positive cells (Fig3.D). This indicates that only RHV1 pseudotyped lentivirus vectors can mediate viral entry in Hepa1-6 cells resulting in reporter gene expression. This data also indicates that surface-expressed RHV1 E1E2 heterodimers are functional.

### Reverse genetics approach did not lead to establishment of an infections clone

An important strategy for HCV antagonizing induction of IFN-mediated innate immunity is cleavage of the retinoic acid-inducible gene I pathway adaptor mitochondrial antiviral-signalling protein (MAVS) by the viral NS3-4A protease(26). Here we used a human hepatoma cell line (Huh-7.5) containing a MAVS cleavage reporter. Briefly, the carboxy-terminal region of MAVS encoding the NS3-4A recognition site and a mitochondrial targeting sequence, has been fused to red fluorescent protein (RFP-MAVS), and a nuclear localization sequence (NLS) was included between the RFP and the MAVS segment. Upon HCV NS3-4A cleavage of the reporter, the RFP translocates to the nucleus(27).

Upon transfection of Huh7-MAVS-RFP cells with a plasmid containing the RHV1 full-length consensus clone, we observed nuclear translocation (Fig. 3E). This confirmed that RHV1 translation occurs in human hepatoma cells and that the RHV1 NS3-4A protease is capable of cleaving human MAVS. RHV1 did not replicate in these cells, however, because the percentage of cells with nuclear translocation decreased and no dsRNA was detected with an anti-dsRNA antibody.

To further asses the ability of the RHV1 and RPGV clones to replicate, we transfected Hepa1-6 and BHK-21 cells with full-length consensus clones harbouring a red fluorescent reporter (mScarlet) and the antibiotic resistance gene blasticidin (BSD) inserted in-between NS5A/NS5B and duplicating the cleavage sequence as previously done for HCV reporters(28). After transfection with the plasmids containing the RHV1 or RPgV sequences we were able to detect mScarlet at 48 hours post transfection. We also observed ds-RNA in a few BHK cells transfected with RHV1 using an anti-dsRNA antibody at 72 hours post transfection (Fig3.F). However, we were not able to select for resistant clones using BSD.

## Discussion

The five rodent hepacivirus IRESs we tested showed different levels of ability to drive translation using in a monocistronic vector across varying cell lines. While the use of a monocistronic vector, utilizing RNA polymerase I, avoids the potential of readthrough in comparison to bicistronic vectors, its drawback is that we weren’t able to directly compare expression levels between cell types due to their difference in susceptibility to transfection. However, the RHV, RHV3 and RHV-rn1 IRESs were either not functional or drove expression at very low levels. This comes as a surprise as the RHV-rn1 virus has already been shown to replicate in both mice and rats.

Both RHV1 and RHV2 drive high levels of translation in murine hepatocytes, MEFs and BHK-21 cells. In human hepatocytes the RHV2 IRES out performed RHV1 which is of interest as they are similar in sequence and therefore likely to have a similar structure. In the case of RHV1, deleting the predicted initial three stem loops abrogates IRES function, suggesting that the full 5’UTR sequence is required to maintain high levels of expression. Also, unlike HCV, where previous studies have shown that the inclusion of 12-30nt of the core protein coding sequence was essential for an efficient IRES activity(29), additional nucleotides from the core protein of RHV1 did not increase transcription levels.

The RPgV 5’UTR has no significant similarity with any known pegivirus but drives high levels of expression in all cell types tested. By deleting specific regions, we were able to show that the initial 126nt of the 5’UTR do not contribute to IRES function and that stem loops IIId/e are essential for maintaining high levels of expression. It would be of significance in the future to confirm the predicted structures of RPgV and RHV1 IRES’s potentially using RNA SHAPE.

RHV1 structural genes (C, E1, E2) expressed from plasmid were shown to be cleaved and yielded proteins of the correct size. Moreover, E1E2 supported transduction of hepatocytes when used to pseudotype lentivirus vectors. Further studies will need to be carried out to find the specific entry receptors, initially blocking CD81 and HCV entry receptors and testing susceptibility of transduction in alternative cell lines, but this initial experiment hints at the hepatotropic potential of RHV1 in mice. This study also confirmed the previous finding that the RHV1 NS3-4A protease is capable of cleaving human MAVS(30).

Our efforts to make an RHV and RPgV replication competent model *in vitro* have so far proved unfruitful. This is of no surprise due to the difficulties in attempting to culture HCV. Both Hepa1-6 and BHK-21 cells failed to yield a clone when selecting for replication with BSD, even when expressing Sec14L2 and ApoE, both essential for high levels of HCV replication(31,32) (not shown). Further cell lines could be tested along with knocking out the innate immune response.

In Summary, this study shows that RHV1/2 and RPgV contain IRESs that are capable of driving high levels of protein synthesis. RHV1 structural genes are cleaved by cellular proteases and can be used to pseudotype lentivirus vectors that are capable of transducing murine hepatocytes.

---

Fig S1: Illustration of plasmid construction as outlined in methods.

