## Supplementary figures and images for "In vitro comparison of the internal ribosomal entry site activity from rodent hepacivirus and pegivirus"

### S1

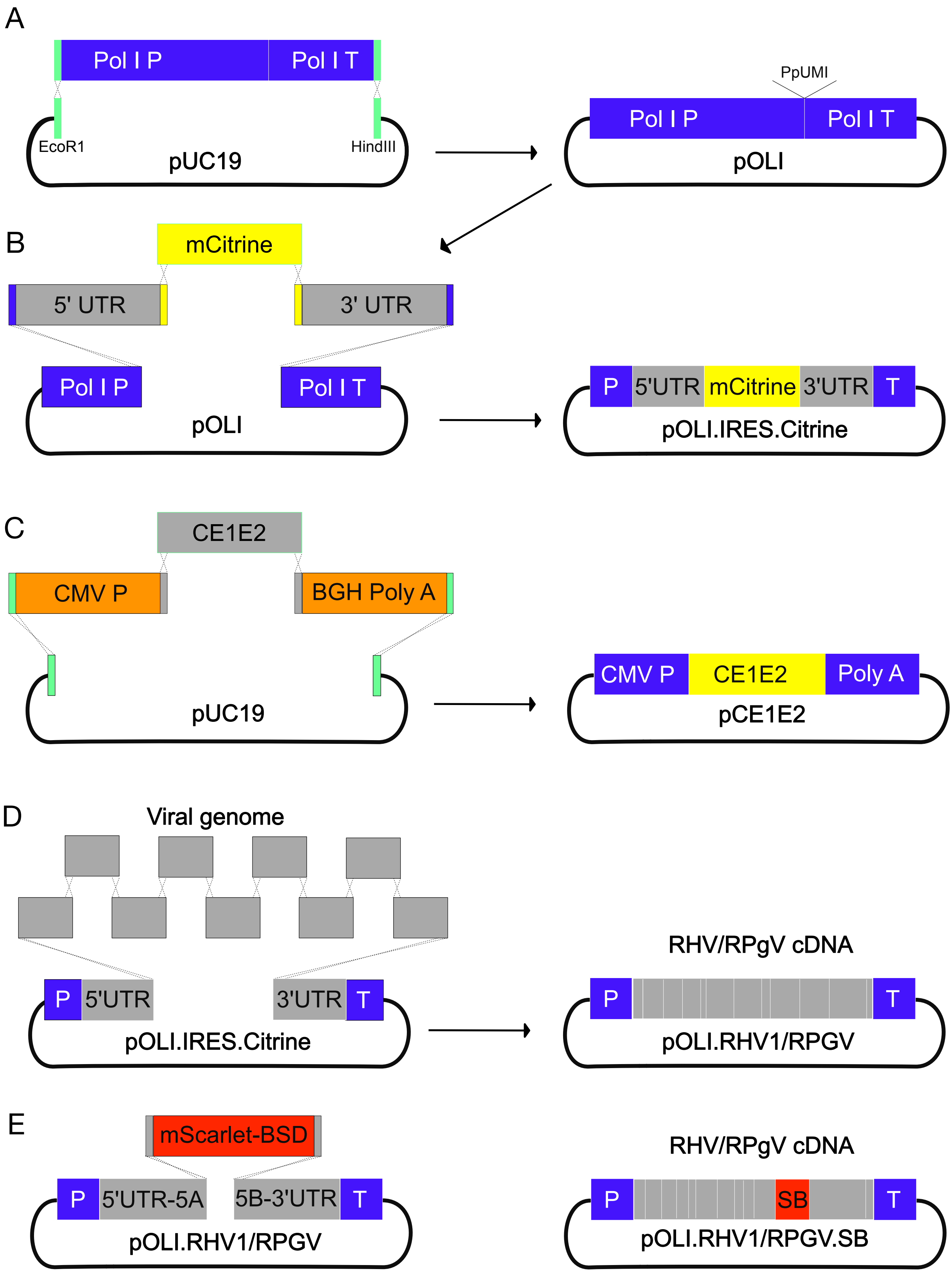
